# Cranial Bone Growth in Isolated Sagittal Craniosynostosis Compared to Normal Growth in the First Six Months of Age

**DOI:** 10.1101/528869

**Authors:** Ezgi Mercan, Richard A. Hopper, A. Murat Maga

## Abstract

**Background:** Sagittal craniosynostosis (SCS), the most common type of premature perinatal cranial suture fusion, results in abnormal head shape that requires extensive surgery to correct. It is important to find objective and repeatable measures of severity and surgical outcome to examine the effect of timing and technique on different SCS surgeries. The purpose of this study was to develop statistical models of infant (0-6 months old) skull growth in both normative and SCS subjects (prior to surgery). Our goal was to apply these models to the assessment of differences between these two groups in overall post-natal growth patterns and sutural growth rates as a first step to develop methods for predictive models of surgical outcome.

**Methods and Findings:** We identified 81 patients with isolated, non-syndromic SCS from Seattle Children’s Craniofacial Center patient database who had a pre-operative CT exam before the age of six months. As a control group, we identified 117 CT exams without any craniofacial abnormalities or bone fractures in the same age group. We first created population-level templates from the CT images of the SCS and normal groups. All CT images from both groups, as well as the canonical templates of both cohorts were annotated with anatomical landmarks, which were used in a growth model that predicted the locations of these landmarks at a given age based on each population. Using the template images and the landmark positions predicted by the growth models, we created 3D meshes for each week of age up to six months for both populations. To analyze the growth patterns at the suture sites, we annotated both templates with additional semi-landmarks equally spaced along the metopic, coronal, sagittal and lambdoidal cranial sutures. By transferring these semi-landmarks to meshes produced from the growth model, we measured the displacement of the bone borders and suture closure rates. We found that the growth at the metopic and coronal sutures were more rapid in the SCS cohort compared to the normal cohort. The antero-posterior displacement of the semi-landmarks indicated a more rapid growth in the sagittal plane in the SCS model compared to the normal model as well.

**Conclusions:** Statistical templates and geometric morphometrics are promising tools for understanding the growth patterns in normal and synostotic populations and to produce objective and reproducible measurements of severity and outcome. Our study is the first of its kind to quantify the bone growth for the first six months of life in both normal and sagittal synostosis patients.

## 1. Introduction

Sagittal craniosynostosis (SCS) is the most common type of premature perinatal cranial suture fusion that results in progressive abnormalities in cranial shape. Although our understanding of the genetics of SCS has increased over the years, (1–4) clinical care of these patients has remained much the same, requiring substantial surgical corrections to address concerns regarding normal brain development due to restricted intracranial volume (ICV) or increased intracranial pressure. Surgeries can be divided into techniques that correct head shape in a single stage from those that require either internal devices or post-operative molding helmet treatment to achieve their goals (5). Common to all procedures is that the infant or young child must be anesthetized for a 1 to 3-hour period for a surgery that results in considerable blood loss (SCH Craniofacial Clinic data). The duration of anesthesia and the volume of blood transfusion during any type craniofacial surgery have been associated with potentially poor neurodevelopmental outcomes in patients later in life (6–8). Since these surgeries are typically performed in the first six months of age, it is important to identify the optimal time to intervene for best surgical outcome while having the least detrimental impact to the patient. (9–13). An increased understanding of the shape changes and suture closure patterns that occur in SCS compared to normal children in the first six months of age would provide valuable information in pursuit of this goal.

### Current challenges in CS outcome research

There are both biological (high-rate of skull and brain growth in infants), and non-biological (choice of surgical technique, age at surgery, socio-economic status) variables that may impact the surgical outcome. Although craniosynostosis cases can be considered common for a structural birth defect, they are still relatively rare conditions (1/1,000 for non-syndromic CS, and less frequent for syndromic cases)(14). These variables, as well as the paucity of large datasets, contribute to the wide range of results and contradicting opinions that exist in the craniosynostosis surgical literature. For example, the important metric of intra-cranial volume (ICV), which CS surgeries are designed to increase, have been reported to be significantly higher (15), lower (16,17) as well as not different in pre-operative synostosis patients compared to controls (18–21). Such variable results can be attributed to low sample sizes, (only one study had 30 cases in any group (20), and some studies had sample sizes as low as 5 per group), to differences in cohort selection (different types of CS might be lumped together), or to methodology. There is also no clear consensus on how to measure the surgical outcome objectively in a longitudinal fashion (22). Long-term studies have indirectly evaluated surgical outcome by comparing the neurodevelopmental outcomes of CS patients who have undergone surgery to non-CS children (23–26). While these studies tend to factor in surgical technique as a variable, they did not always control for the initial severity of the condition and the magnitude of surgical correction. Even when the severity was measured, the metrics used were simplistic, and did not represent the full spectrum of disorder (15,27–29). One study that used the facial appearance of children with CS who underwent surgery found that lay-people do not perceived them as ‘normal’ compared to unaffected children, even after surgery (30).

In summary, there is an acute need to find objective and repeatable measures of severity and surgical outcomes in SCS treatment and planning. As the first step towards this goal, we developed statistical models of infant (0-6 months old) skull growth both in normative and SCS patients and applied them to the assessment of differences in both overall post-natal growth patterns and sutural growth rates. By studying the shape and suture changes in SCS patients during this critical period, we can better understand the implications of operative intervention at a given age on final surgical outcome.

## 2. Materials and Methods

The study was reviewed and approved by the Institutional Review Board at Seattle Children’s Hospital (STUDY00000167, STUDY00000424).

### 2.1 Dataset and Statistical Analysis

There were three distinct steps in our full analysis: Estimation of canonical templates for both normal and SCS cohorts; statistical modeling and visualization of cranial growth; and estimation of bone growth and suture closure rates. Below we provide the specifics of each step. A summary flowchart is also provided as a supplement figure (**S1 Figure).**

#### 2.1.1 Normal Cohort

We retrospectively reviewed all patients under the age of six months with a head CT exam performed at Seattle Children’s Hospital between 2004 and 2017. The patients with skull abnormalities, bone fractures, head/face/ear deformities and congenital conditions (including growth restrictions) were excluded after a review of radiology reports and clinical histories. For the remaining patients, CT scans were 3D rendered in bone window and visually inspected to discard any low-quality or substantially incomplete field of view of cranium.

Our final control cohort included 117 CT exams. Of the 117 exams, 79 had a complete cranium. In the remaining 38 CT exams maxilla was cut below the nasal spine, yet all the pertinent anatomical landmarks were identifiable. All 117 exams were included in the control population. The most common reasons for a head CT exam with no abnormal findings were head injury, seizure and scalp swelling.

#### 2.1.2 Sagittal Craniosynostosis Cohort

Seattle Children’s Craniofacial Center maintains an independent database of patients seen in the clinic. We reviewed this database to identify patients under the age of six months with an isolated sagittal craniosynostosis diagnosis with a pre-operative CT scan performed at our institution between 2004 and 2017. We visually inspected CT scans by rendering them in bone window to discard any low-quality or incomplete scans. Since pre-operative CT scans were performed for surgical planning or diagnostic purposes, almost all scans provided a high-quality and complete image of the cranium. Our sagittal craniosynostosis cohort consisted of CT scans from 81 patients.

### 2.2 Anatomical Template Building

We built both a normal and a SCS population-level anatomical template for infants 0-6 months of age. Because we expected that major changes in the normal skull form would be driven entirely by biological growth, we wanted the normative canonical template to be unbiased with respect to age, as well as sex. Therefore, we selected 34 individuals from our normal cohort, such that CT exams from a male and a female patient approximately represent one week of age between 0-6m.

For the SCS cohort, we had no such *a priori* expectation, and qualitatively the phenotypic variability of skull forms appeared to be independent of the age. Moreover, SCS tends to be more common in males than females (with an approximate ratio of 4:1 (14)), making it harder to identify a sub-cohort that is balanced both for age and sex. Therefore, we opted to use all of our SCS cohort to generate SCS.

All CT images in this study were manually edited to remove mandible and other non-cranial bones and artifacts (e.g., tubes, pacifiers) using Segment Editor in 3D-Slicer (31). A global threshold was used to extract only the bony cranium from CT exams, and all samples were rigidly aligned to a reference sample to remove positional differences. At this step, all exams were mirrored along the midsagittal plane, and these mirror images included in the anatomical template building process to generate bilaterally symmetric templates.

We used the multivariate template construction pipeline that is provided with the Advance Normalization Tools (ANTs) image registration and analysis ecosystem (32,33). ANTs template-building script estimates the best average shape that represents the entire population by iteratively calculating an average image and registering each sample to the new average. We used the default registration parameters: the geodesic diffeomorphism transformation of symmetric normalization (SyN) and the similarity metric of cross-correlation (33). We found that six iterations were sufficient to construct an anatomically refined template, and after the fifth iteration, the changes in the template were negligible. The resultant template reviewed by a morphologist (AMM) and a craniofacial surgeon (RH) for anatomical accuracy.

### 2.3 Landmarks and Semi-landmarks

We rendered all exams in bone window and annotated 38 anatomical landmarks (See supplementary material **S2 Appendix**) using 3D-Slicer program (31). A subset of five control and five sagittal craniosynostosis images were independently annotated by two observers (EM and AMM) to measure inter-rater variability and the annotations were repeated after a wash-out period of one month to measure intra-rater variability. After obtaining low intra- and inter-rater variances for all landmark points, all CT scans in both cohorts were annotated (by EM). **Fig 1** illustrates the landmarks on the normative template. The cranial landmarks were annotated on all CT scans and used to model growth in two populations.

**Fig 1:**
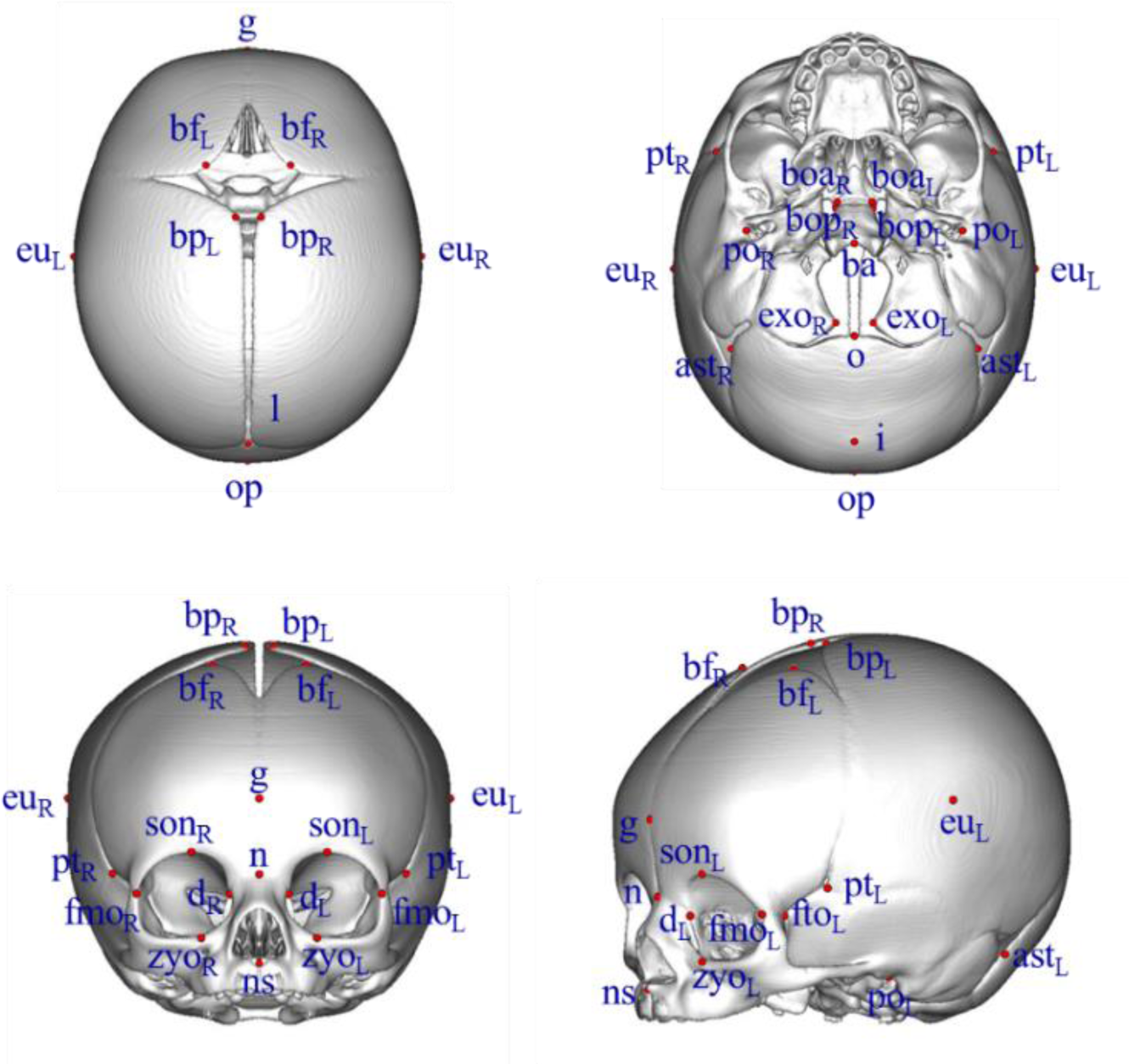
38 cranial landmarks used in the study (the landmark vertex, “v”, was removed in control samples since it does not fall onto a bony structure in young infants with a patent sagittal suture)

While cranial landmarks represented the standard anthropometric features used to assess overall skull shape and size in literature, they are not sufficient to measure precisely the growth and size changes along the major cranial sutures. We therefore annotated the major cranial sutures (metopic, coronal, sagittal and lambdoidal) on both templates with semi-landmarks equally spaced within regions as shown in **Fig 2**. Both templates had a total of 80 matched semi-landmark points to allow direct comparison. Since the normal and SCS templates were symmetrical, only the left side of the cranium was annotated with the semi-landmarks. The two annotated templates were automatically transferred to the meshes produced with the landmark-based growth model. The details of semi-landmark annotation are in Appendix A.

**Fig 2:**
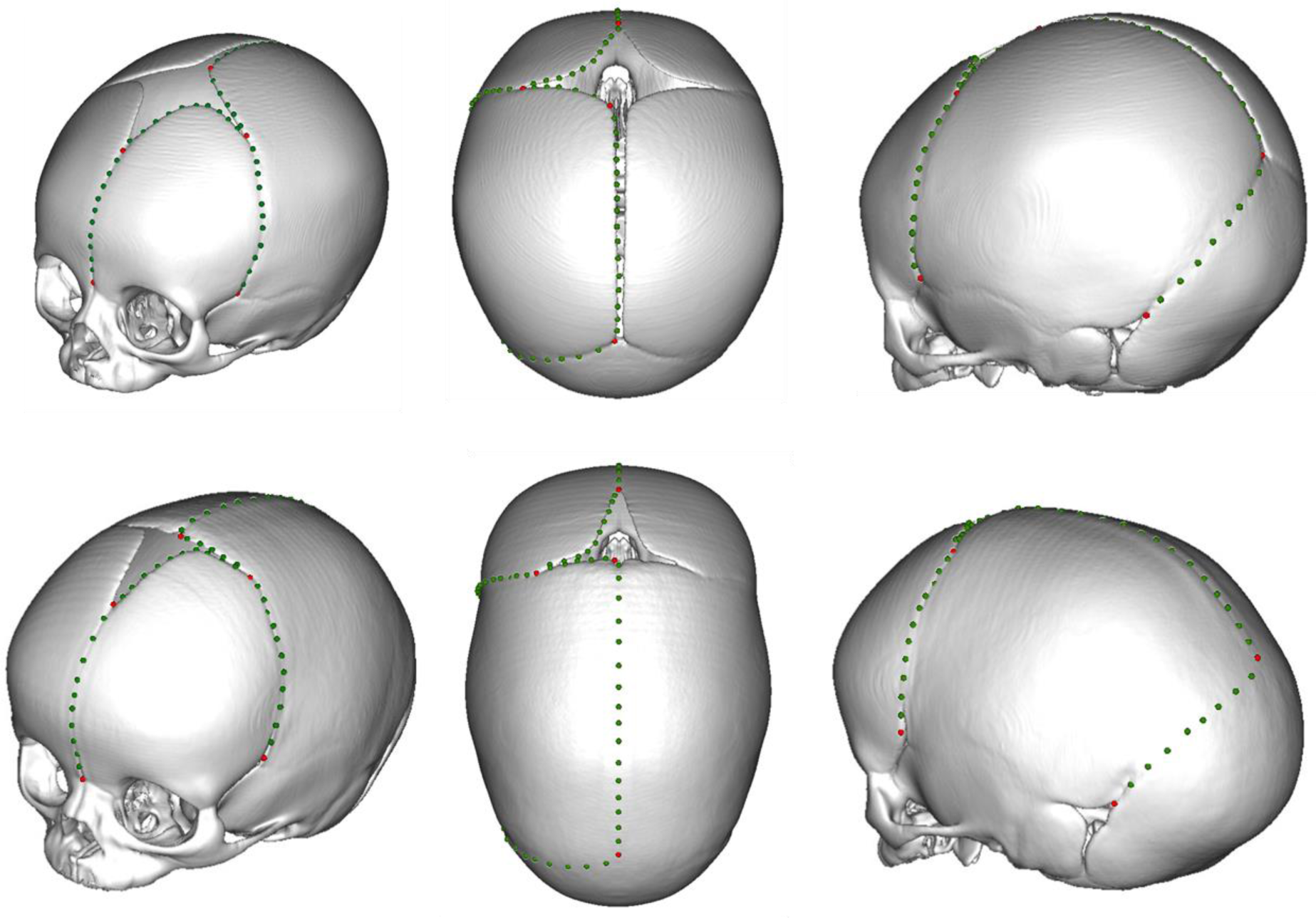
Semi-landmarks placed along the metopic, coronal, sagittal and lambdoidal sutures to estimate the suture closure rates in control and sagittal craniosynostosis populations. Red points correspond to the borders of different regions based on sutures and fontanelles. Top row: normal template, bottom row: sagittal craniosynostosis,

### 2.4 Growth Models

We modeled skull growth using the cranial landmarks and patient age. The models predicted the locations of the cranial landmarks over the age range 0 to 6 months. The normal growth model included 117 samples (a total of 234 with mirrored samples) and the sagittal craniosynostosis growth model included 81 samples (a total of 162 with mirrored samples). Since the young infants in the control dataset have a patent sagittal suture, the vertex point (the highest point of the head) was not always on a bony structure and was thus excluded from the normal growth model. Similarly, the inion had a high variance in the sagittal craniosynostosis population and was excluded from sagittal craniosynostosis growth model. Cranial landmarks were aligned using Generalized Procrustes analysis (GPA) with the geometric morphometrics package Morpho in R. (34,35). Because we wanted to model cranial growth in physical space, the uniform scaling step of the GPA analysis was skipped

Principle component analysis (PCA) is a workhorse of the geometric morphometric methods (36). It converts a set of observations (landmark coordinates) into uncorrelated latent variables sorted in the order of decreasing variance. PCA is commonly used for feature selection by eliminating the components with small variances. In our models, we used the first 20 principle components (PCs), which explained ∼90% of the variation while reducing the number of variables from 111 to 20. PCA also provides uncorrelated variables that can be modeled independently in contrast to the “raw” coordinate values that are highly correlated in cranial growth. Several regression models (linear, local, polynomial) were evaluated in a leave-one-out-cross-validation (LOOCV) setting (See supplementary material **S3**) and the one with the least error and the least complexity, linear regression, was chosen. All statistical modeling was conducted in R.

### 2.5 Bone Growth and Suture Closure

Using the growth models, we estimated landmark positions for each week of age up to 6 months for both the control population and the sagittal craniosynostosis population. Landmark positions provide control points that allow a surface mesh to be interpolated to estimate the shape and size changes in growing skulls. We extracted surface meshes from the volumetric template images using the marching cubes algorithm in VTK library (37,38) and warped the template meshes to the target landmark points at different ages using the thin plate splines (TPS) method from the package Morpho in R (35,36).

The semi-landmarks from the templates were transferred to the meshes at different ages produced by TPS. Since the orientation of the mesh determines the growth direction, meshes were aligned to Frankfort Horizontal using the landmarks porion and zygoorbitale and translated to have the basion as the origin. Finally, the displacement of each semi-landmark was calculated between consecutive weeks and averaged to obtain a growth rate.

## 3. Results

### 3.1 Principal Component Analysis of the Landmarks

To better understand the shape changes associated with each PC, we visualized the eigenvectors associated with each PC. By changing the PC scores between two standard deviations around the mean of their respective populations, we created synthetic meshes that illustrate the skull growth as captured by each PC.

For our study population, PC1 (visualized in **Fig 3**) explained 35% of the variation in control cohort and 30% of the variation in sagittal craniosynostosis cohort. In both populations, PC1 modeled the overall skull size and the closure of the anterior fontanelle (as noted by red arrows in **Fig 3**). Because change in skull size is strongly correlated with age in young infants, PC1 also strongly correlated with the age in both populations (*R*^2^ = 0.68 for normal cohort and *R*^2^ = 0.59 for sagittal craniosynostosis cohort) (See supplementary material **S4 Figure**).

**Fig 3:**
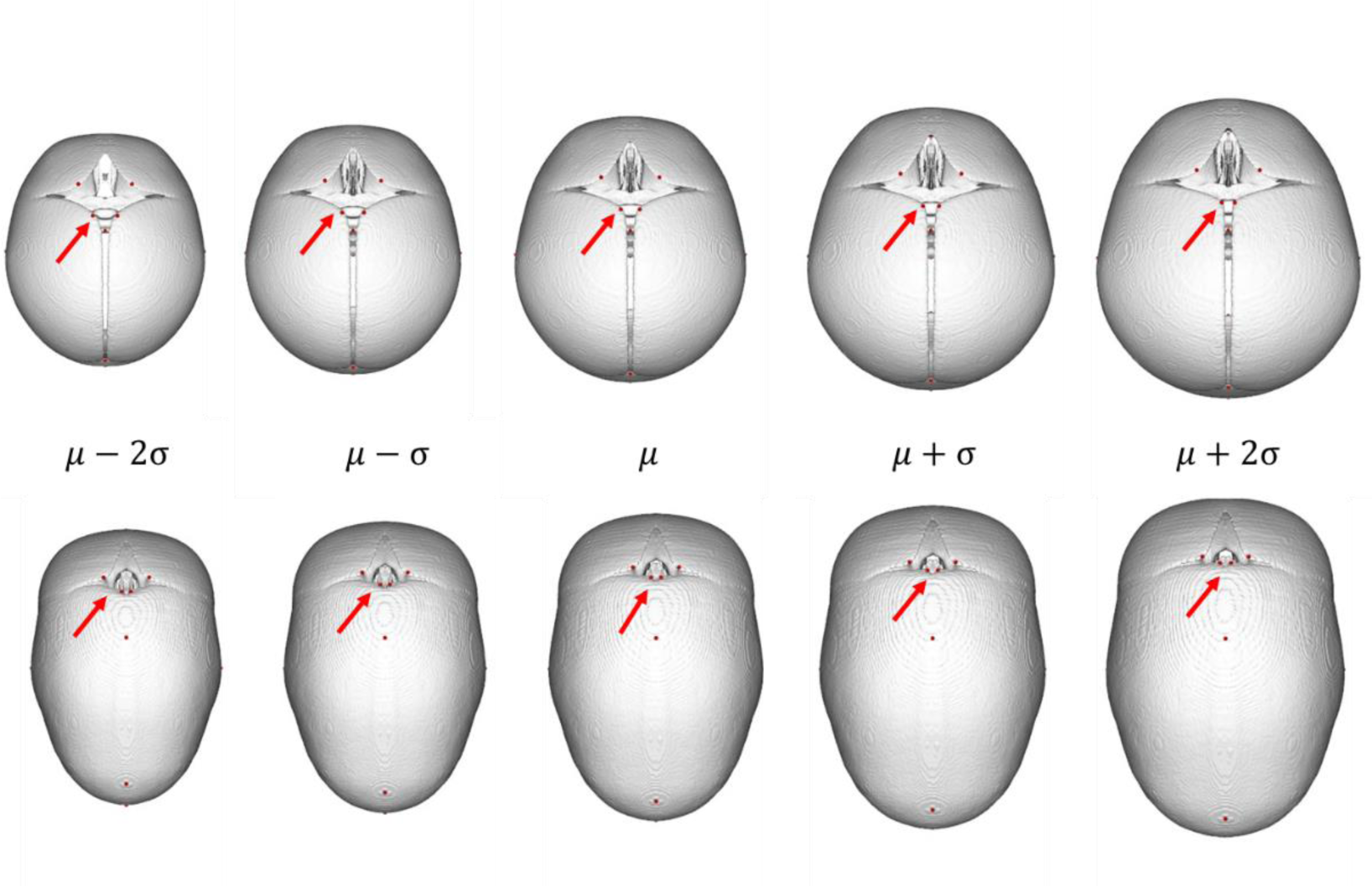
The shape variation captured by the first principle component. The top row shows the reconstructed normative template and the bottom row shows the reconstructed sagittal craniosynostosis template using the first PC scores in two standard deviations around the mean of their respective populations.

### 3.2 Bone Growth

We quantified the bone growth and suture closure rates by measuring the displacements of semi-landmarks that were transferred to surface meshes for each week of age from 11 days to 179 days. Although control cohort had younger (3 days old) and older (183 days old) patients, we did not have any samples to model the sagittal craniosynostosis growth for these extreme ages hence we limited our growth analysis to a slightly narrower age range for the sagittal craniosynostosis cohort (11 to 179 days old). The **Fig 4** shows surface meshes and semi-landmarks for five samples generated by the growth models for both cohorts. We analyzed the growth in three major axes: horizontal (mediolateral), anteroposterior and craniocaudal. To further simplify the interpretation, we divided the semi-landmarks into seven regions as described in **Table 2.**

**Table 1:**
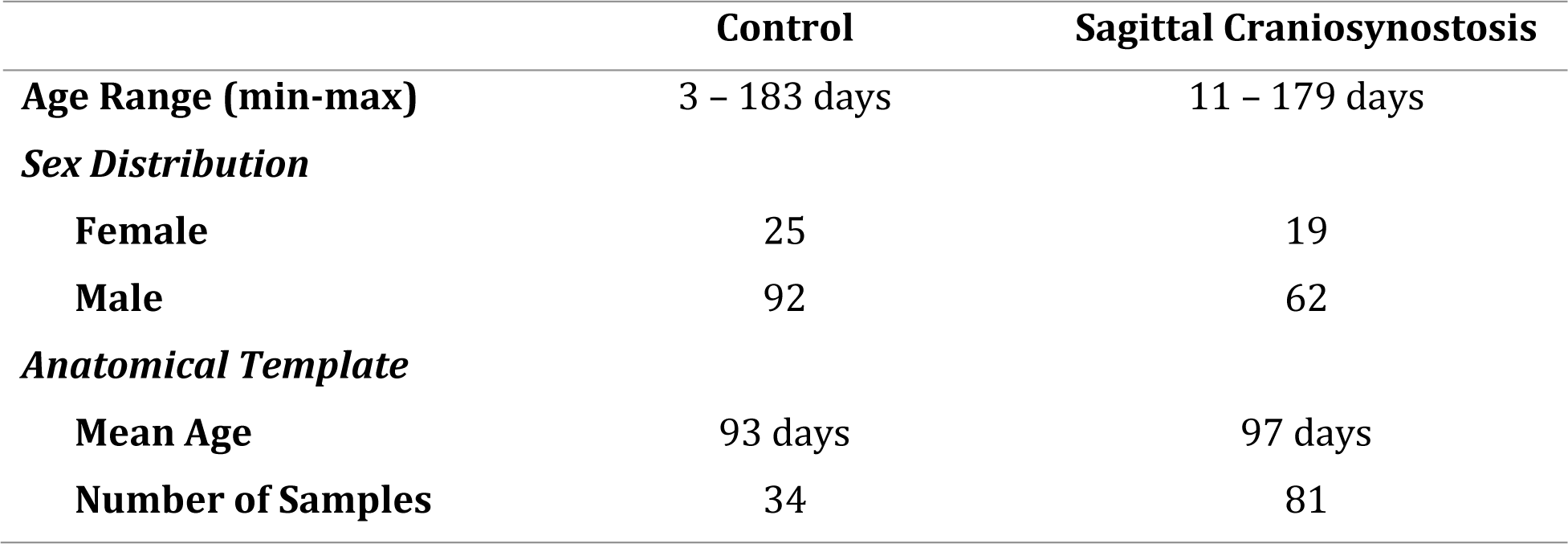
Demographics-Sagittal craniosynostosis and control cohorts.

|  | Control | Sagittal Craniosynostosis |
| --- | --- | --- |
| **Age Range (min-max)** | 3 – 183 days | 11 – 179 days |
| ***Sex Distribution*** |  |  |
| **Female** | 25 | 19 |
| **Male** | 92 | 62 |
| ***Anatomical Template*** |  |  |
| **Mean Age** | 93 days | 97 days |
| **Number of Samples** | 34 | 81 |

**Table 2:** Semi-landmark regions along the metopic, coronal, sagittal and lambdoidal sutures.

| Region | Description |
| --- | --- |
| **Region 1** | The metopic suture from the nasion to the anterior-most point of anterior fontanelle at the mid-sagittal plane. |
| **Region 2** | Anterior border of the anterior fontanelle on the frontal bone from the metopic suture to the coronal suture |
| **Region 3-4** | The coronal suture between parietal and frontal bones from the from the lateral apex of the anterior fontanelle towards the pterion constitutes region III while the reverse direction from the pterion towards the anterior fontanelle constitutes region IV. |
| **Region 5** | The posterior border of the anterior fontanelle on the parietal bone from the coronal to the sagittal suture. |
| **Region 6** | The sagittal suture (fused on the sagittal synostosis template and unfused on the normal template) between two parietal bones from the posterior-most point of the anterior fontanelle on the mid-sagittal plane to the landmark lambda. |
| **Region 7** | The lambdoidal suture from the landmark lambda to the edge of the mastoid fontanelle. |

**Fig 4:**
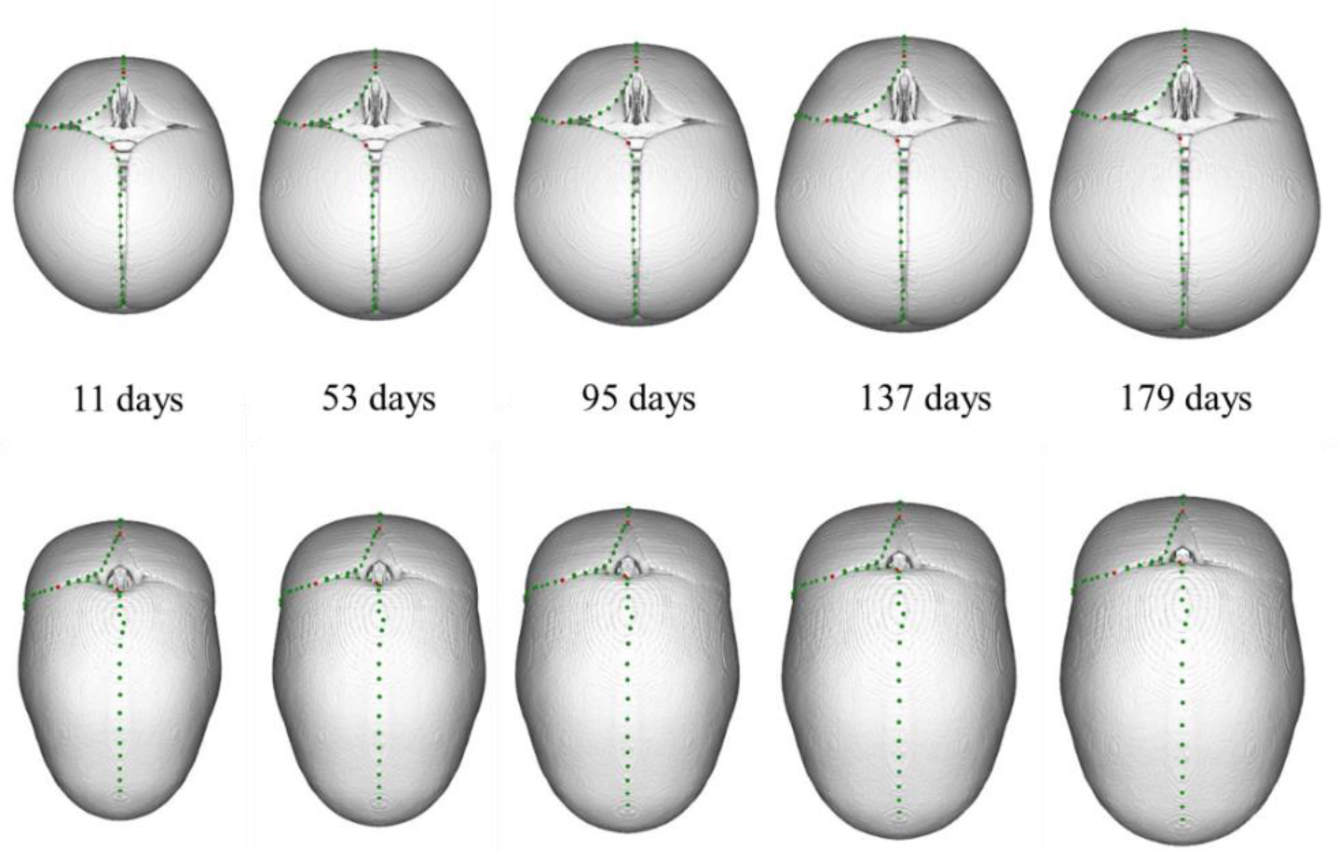
Visualizations of predicted growth from the statistical model along selected time points in normal (top row) and craniosynostosis populations (bottom row). An animation of continuous visualization is provided as a supplementary file (S5).

#### 3.2.1 Mediolateral Growth

Fig **5**A shows the normal growth vs. growth in sagittal craniosynostosis in the mediolateral axis. Y-axis shows the displacement (mm/week) at semi-landmark positions where positive values indicate medial movement (can be interpreted as suture closure) and negative values indicate lateral movement (associated with overall growth of the skull). Displacement along the metopic suture (region I) and the sagittal suture (region VI) of both normal and SCS models was negligible. The displacement of frontal bone suture landmarks along the anterior border of the anterior fontanelle (region II) was in the lateral direction for the normal model. This movement away from the mid-sagittal plane is indicative of an overall regional growth in the normal model that was faster than the fontanelle closure (medial displacement) at the metopic and coronal sutures. In comparison, the overall medial movement of the corresponding region II suture landmarks in the sagittal craniosynostosis model indicates that fontanelle closure was more rapid than overall lateral cranial growth. The displacement in regions III and IV, which corresponds to movement along the coronal suture, was in the lateral direction for both models, indicating lateral cranial growth, but it was smaller for the SCS model, indicating a restricted horizontal growth relative to normals. Parietal bone landmarks in region V, along the posterior border of the fontanelle moved in a medial direction, particularly as they approached the sagittal suture, indicating parietal bone growth towards the anterior fontanelle in both groups. This medial displacement rate was faster in the SCS model.

**Fig 5:**
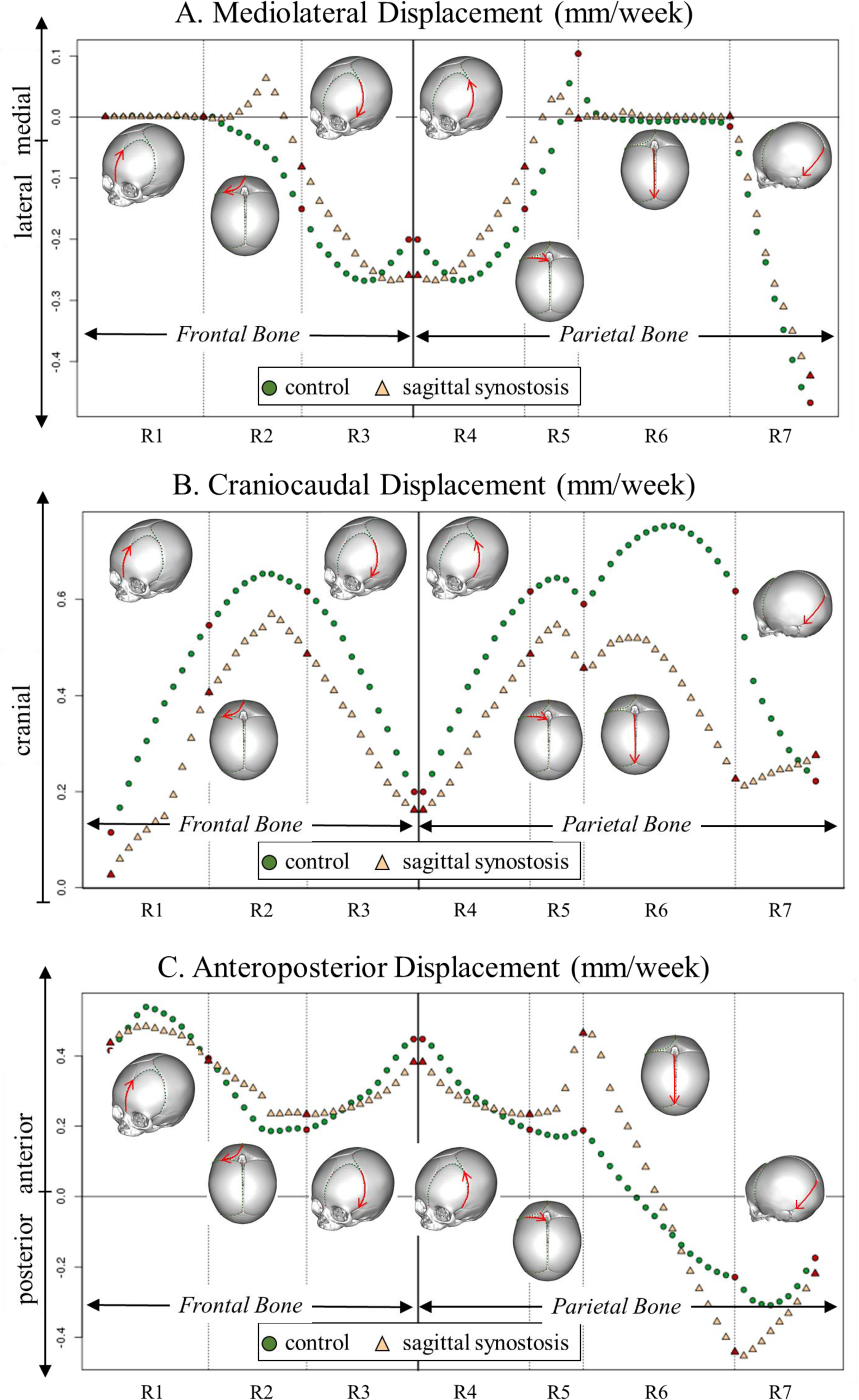
Normal growth vs. growth in sagittal craniosynostosis. A. Mediolateral displacement: positive values indicate medial movement (can be interpreted as suture closure) and negative values indicate lateral movement (mostly associated with the overall growth of the skull). **B.** Craniocaudal displacement: positive values indicate a cranial movement. **C.** Anteroposterior displacement: positive values indicate anterior, and negative values indicate posterior movement with respect to basion.

#### 3.2.2 Craniocaudal Growth

**B** shows the displacement of semi-landmarks in the longitudinal axis for normal and sagittal craniosynostosis growth models. In all regions, the displacement was in the cranial direction due to the alignment of the model with basion at the origin. There was no vertical compression of the skull with age. The cranial displacement indicates overall skull growth that was faster in the normal model compared to the SCS model. The slower vertical or cranio-caudal growth of the sagittal craniosynostosis model indicated relative growth restriction in the longitudinal axis. Interestingly, the pace of growth was considerably slower at the posterior cranium (region VI) for the sagittal craniosynostosis model, which likely contributes to the decreased vertical posterior cranial height and occipital protuberance phenotype commonly observed at the time of surgery in these patients.

#### 3.2.3 Anteroposterior Growth

**Fig 5C** illustrates the displacement of semi-landmarks in the anteroposterior axis for normal and SCS growth models. The positive values in the y-axis indicate anterior movement whereas the negative values indicate posterior movement. The consistent anterior displacement of semi-landmarks in regions I through IV indicate overall anterior skull growth faster than the bone growth of the frontal bones into the anterior fontanelle. The positive displacement of semi-landmarks in region V indicates greater growth of the parietal bone into the anterior fontanelle compared to posterior displacement from overall cranial growth. This anterior displacement rate was faster for the SCS model, indicating more rapid closure of the anterior fontanelle to normal model from this posterior border. The anterior movement of the semi-landmark at the border of region V and VI, the point where the sagittal suture meets the coronal sutures, was much faster in the SCS model than the normal model. These semi-landmarks were posterior to the basion (the point of origin for alignment) at the second half of the region VI, and they move posteriorly for both models, indicating skull growth. The posterior growth was faster for sagittal craniosynostosis model which contributes to the scaphocephalic elongated skull shape of the sagittal craniosynostosis patient.

### 3.3 Frontal Bone Growth

We compared the semi-landmark movements in R1 and R3, two borders of the frontal bone along the metopic and coronal sutures. Semi-landmarks in both regions moved anteriorly but mode rapidly in R1, suggesting an increase in the distance between the metopic and coronal sutures and an overall anteroposterior growth in the frontal bone in both models (**Fig 5C**). Similarly, the semi-landmarks at the metopic suture (R1) did not move mediolaterally in both models but the semi-landmarks at the coronal sutures (R2) moved laterally in both models (**Fig 5A**). This indicated a mediolateral growth of the frontal bone but at a much rapid pace in the normal model.

### 3.4 Parietal Bone Growth

The semi-landmarks in R4, R6 and R7 marked the borders of the parietal bone along the coronal, sagittal and lambdoidal sutures. The coronal suture semi-landmarks (R4) and lambdoidal suture semi-landmarks (R7) moved laterally in both models while the sagittal suture semi-landmarks did not move mediolaterally, indicating mediolateral growth of the parietal bone in both models (**Fig 5A**). Lateral movement of the semi-landmarks in R4 and R7 was faster in normal model indicating a restriction in mediolateral growth of the parietal bone in SCS model. All semi-landmarks surrounding parietal bone (R4-R7) moved superiorly in both models (**Fig 5B**). The superior movement was faster in normal models, again, indicating a superoinferior growth restriction of the parietal bone in SCS model. Finally, the coronal suture semi-landmarks (R4) moved anteriorly while the lambdoidal suture semi-landmarks moved posteriorly in both models (**Fig 5C**). The movement was faster in both direction for the SCS model, indicating a faster anteroposterior movement of the parietal bone compared to normal model which contributes to the scaphocephalic elongated skull shape of the sagittal craniosynostosis patient.

### 3.5 Closure Pattern of the Anterior Fontanelle

The symmetry of the models allowed an assessment of the relative movement of the frontal and parietal bone plates in the region of the anterior fontanelle. Lateral displacement of the suture landmarks would indicate that horizontal growth perpendicular to the metopic and sagittal sutures was greater than the growth of the frontal and parietal bone plates into the anterior fontanelle. In comparison, medial displacement would indicate that anterior fontanelle closure was faster than overall cranial horizontal growth. Leveraging the bilateral symmetry of the model, we reflected the semi-landmarks across the mid-sagittal plane and measured distances between the right-left pairs of semi-landmarks. Using the growth model, we calculated the average weekly change in the distance between bilateral semi-landmarks as shown in **Fig 6**. In normal growth, the overall width of the anterior fontanelle increased at a rate up to 0.3mm/week at the point where it met the coronal, but it decreased in width at the posterior border as it approached the sagittal suture. This would indicate the fontanelle closure was primarily from a medial in-growth of the opposing parietal bone plates. In comparison, in the SCS model, the width of the anterior fontanelle decreased at both its anterior (metopic) and posterior (sagittal) apices. This would indicate that compared to normal fontanelle closure, in SCS there is a greater compensatory ingrowth of the frontal bones, resulting in a circumferential rather than a posterior to anterior closure observed in the normal model.

**Fig 6:**
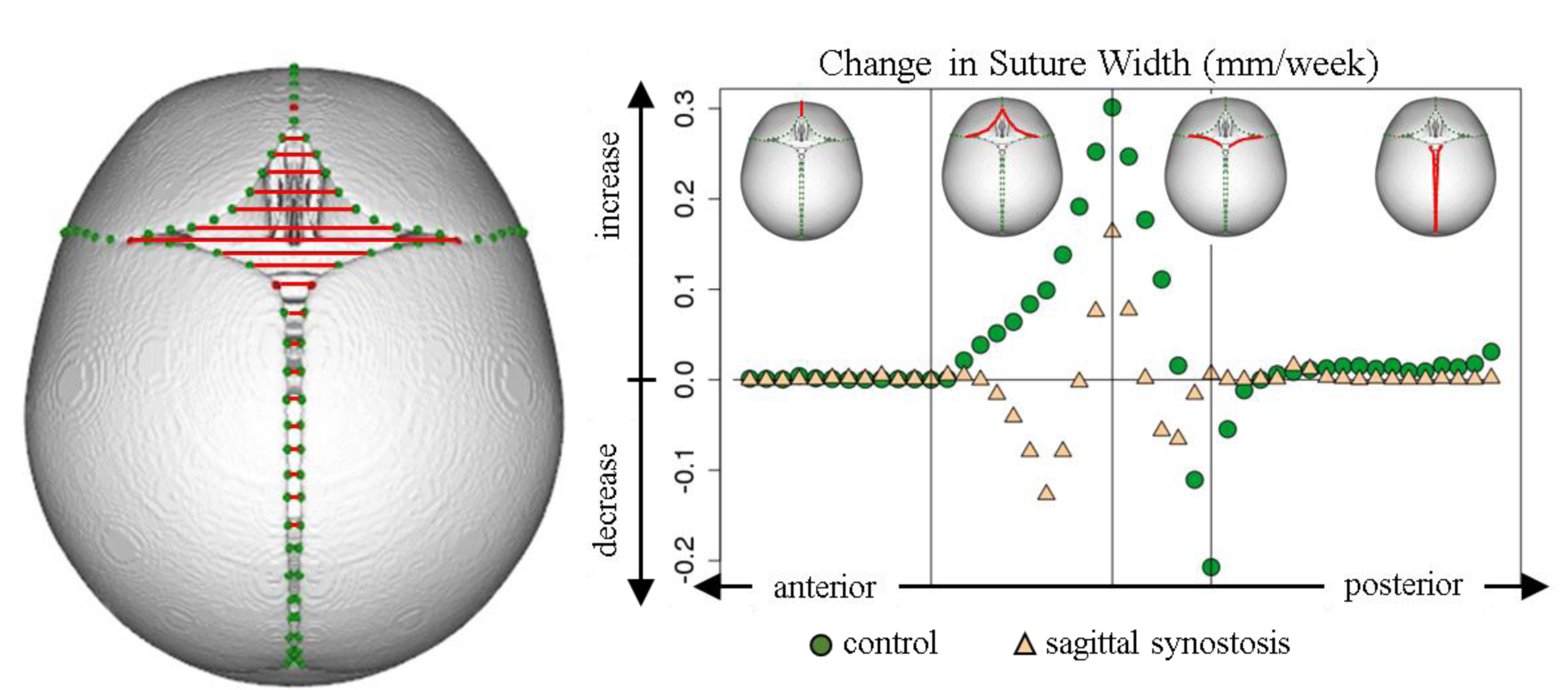
Average change in the distance between bilateral pairs of semi-landmarks.

A similar analysis was conducted by calculating the distances between pairs of landmarks on opposing anterior-posterior sides of the anterior fontanelle. **Fig 7** shows the average change in the distance between the semi-landmarks in a sagittal plane. The distance slightly increases in the normal growth model likely secondary to overall skull growth. However, for the SCS growth model, the anterior-posterior distances decreased. This indicated that the closure of the anterior fontanelle in this direction was greater than the overall parallel growth of the skull. The pace of the fontanelle closure was greatest towards the mid-sagittal symmetry plane.

**Fig 7:**
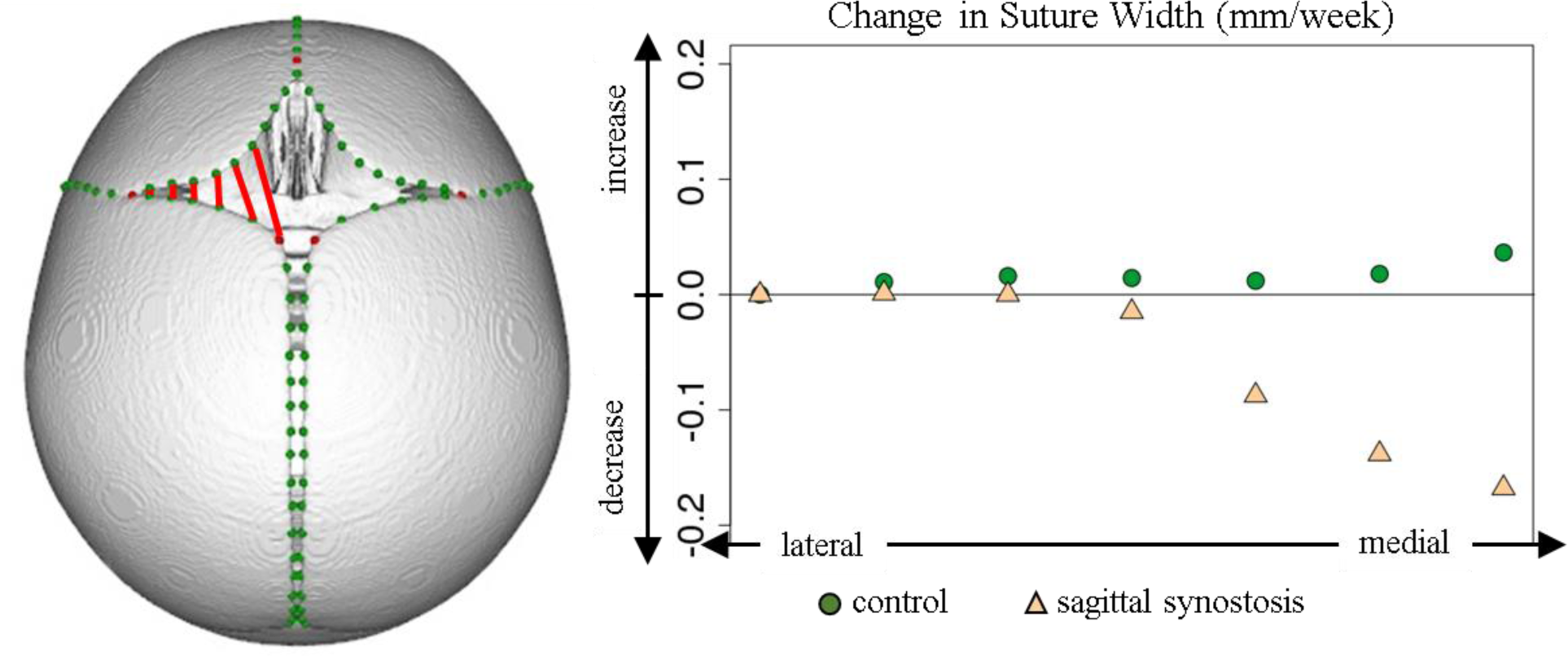
Average change in the coronal suture width along seven semi-landmark pairs.

## 4. Discussion

During the first months after the birth, the infant skull goes through a transformation that is unparalleled at any other point in human life: a newborn cranium is approximately 25% of its adult size and it doubles in size by the first six months (39). This rapid brain development and intracranial volume expansion cause an infant’s skull to expand through the patent sutures between the major cranial bones. The crucial role of major cranial sutures in skull shape and growth is accepted and molecular mechanisms underlying the bone differentiation at the suture sites are being identified by ongoing research (40).

### The Virchow Law and Bone-Plate Theory

Despite the advances made in animal models and molecular experiments, the characteristic shape deformities associated with the sagittal synostosis is still explained by the Virchow principle of 1851 (41) which states that “the growth is restricted in a plane perpendicular to the fused suture, and enhanced in the corresponding parallel plane”. We did observe a more rapid pace of growth in the anteroposterior direction in the SCS model compared to normal which is consistent with this principle. Virchow’s principle does explain the scaphocephalic elongated and narrow cranial shape associated with premature fusion of the sagittal suture, but it does not explain the subtler phenotypic changes such as occipital constriction, decreased posterior vertical height, and pterion construction. In 1991, Delashaw et al observed that asymmetrical bone deposition occurred mainly at perimeter sutures with growth directed away from the fused bone plate. They also stated “As new radiologic techniques develop to define the configuration of the skull in intricate detail, a skull pattern of growth explaining the pathogenesis of all deformities created by premature fusion of a cranial vault suture may become apparent.” We also found a higher rate of bone growth at the coronal sutures around the anterior fontanelle in sagittal craniosynostosis model compared to normal growth model.

Our preliminary analysis with the PCA showed that the first principle component was correlated with the skull size and the patient age. In our population, PC1, the dominant direction of change, also showed a consistent pattern of parietal bone growth around the anterior fontanelle and along the sagittal suture in both cohorts. By annotating the ends of the sagittal suture on parietal bones, namely bregma (bp) and lambda (l) landmarks (See Fig **1** and supplementary material **S2 Appendix**), we were able to capture this dominant pattern across the population.

### Premature Closure of Uninvolved Sutures

The bone growth around unfused metopic and coronal sutures was faster in sagittal craniosynostosis cohort compared to normal cohort. This explains the observation that older kids with sagittal synostosis present with smaller anterior fontanelle. An interesting notion to consider is the phenomenon of delayed synostosis of uninvolved sutures or even pansynostosis (premature fusion of all sutures) after surgical intervention (11,13,42,43). With our findings indicating a tendency to early closure at metopic and coronal sutures in sagittal synostosis cohort, it is worthwhile to investigate the effects of the surgery and the synostosis on the uninvolved sutures separately.

### Statistical Growth Models vs Finite Element Analysis

We modeled the calvarial bone growth during the first six months of life in infants with isolated sagittal craniosynostosis and infants without any craniofacial abnormalities using a cross-sectional population approach in a data-driven way. We were able to create two growth models that confirm the well-established observations, as well as reveal novel findings about the bone growth at the suture sites.

Human skull growth had also been modeled as a mechanical process that is driven by the expansion of the brain and consequently neurocranium using finite-element (FE) methods (44–46). This approach focuses on material properties of the calvarial bones and sutures while our approach relies on the data that CT exams provide. Furthermore, it models the bones as they are in the template used and does not incorporate the bone growth at the suture sites. The driving force in FE models is the brain growth as a physical force determined via in-vivo experiments. The population-level variance in shape and growth is not part of FE analysis while it is an important piece of data-driven statistical models. FE approach allows an analysis of extreme details on one sample while our statistical models enables an analysis at the population-level.

### Manually annotated Landmarks vs. Dense correspondence

We used cranial landmark positions as the response of our growth models based on age. We included landmarks around the anterior fontanelle, such as four bregma points on the corners of frontal and parietal bones, to better model the overall bone growth along the patent sutures. While admittedly sparse, selected cranial landmarks provided us with a sufficient topographic coverage of the skull surface to model the overall cranial growth. This approach can be expanded to use dense correspondence between 3D models of skull. The challenge, however, is that infants, and especially non-craniosynostosis cases, display a wide range of variability in suture shape and closure. This is a challenging case for the automated registration algorithms to find the optimum correspondence without the expert guidance. In this study, we established a baseline growth based on expert knowledge, which we plan on to refine using a combination of anatomical landmarks and semi-landmarks automatically detected on all surfaces which could provide a denser correspondence among samples and help us capture subtle differences in surface areas that are not annotated in the current study.

Frontal bossing and occipital protuberance are common deformities associated with SCS and believed to change over the first six months of age with rapid growth of the brain. Our growth models were based on anatomical landmarks that do not cover the regions of the cranial bones that exhibit these deformities. For this reason, we were unable to capture the shape changes that occur on the bone surfaces without anatomical landmarks. We plan to develop a landmark-free growth model based on dense correspondences between samples. Such models will allow us to interpret growth at sutures and on bone surfaces.

### Future Directions

Our study is the first of its kind to model and compare the skull growth of large normative (N=117) and sagittal craniosynostosis (N=81) populations. It is of clinical importance to establish normative templates and growth models that can be used to evaluate the degree of deformity and the surgical outcomes. We plan to extend our study to other isolated craniosynostoses (such as metopic and unicoronal craniosynostosis) and older age groups. Comparing skull growth and bone deposition patterns at the sutures can provide valuable insight into understanding the etiology of single suture craniosynostosis and guide the surgical treatment.

## 5. Conclusions

Sagittal craniosynostosis is the most common single-suture isolated form of craniosynostosis. The surgical intervention is usually planned in the first year of life, most commonly between 3 to 6 months of age. The brain development and skull growth are at their fastest rate during this period, yet the literature lacks studies with large sample sizes focusing on the shape and growth of the infant skull. In this study, we first generated two anatomical templates, a normative and a sagittal craniosynostosis template, to represent the two distinct populations. Then, using anatomical and geometric landmarks, we modeled the growth of the two populations from birth to six months. Finally, with the help of additional semi-landmarks along the sutures, we compared the bone growth rates between two populations.

We found that the dominant growth pattern in both cases is the overall increase in the skull size and the decrease in the size of the anterior fontanelle in the case of sagittal craniosynostosis population. Our measurements showed that the sagittal craniosynostosis model grows in the anteroposterior direction faster than the control cohort. Additionally, the coronal and metopic sutures close faster in the sagittal craniosynostosis model compared to normative growth.

## Supporting information

Appendix: Landmarks and semi-landmarks

Appendix: Statistical models compared

supplemental animation

---

S1 Figure: Summary flowchart

S2 Appendix: Landmarks and Semi-landmarks

S3 Appendix: Comparison of Growth Models

S4 Figure: PC1 scores are highly correlated with age at the time of exam.

S5 Figure: Growth animations for normal and SCS models.

