## Supplementary figures and images for "Cranial Bone Growth in Isolated Sagittal Craniosynostosis Compared to Normal Growth in the First Six Months of Age"

### supplemental animation

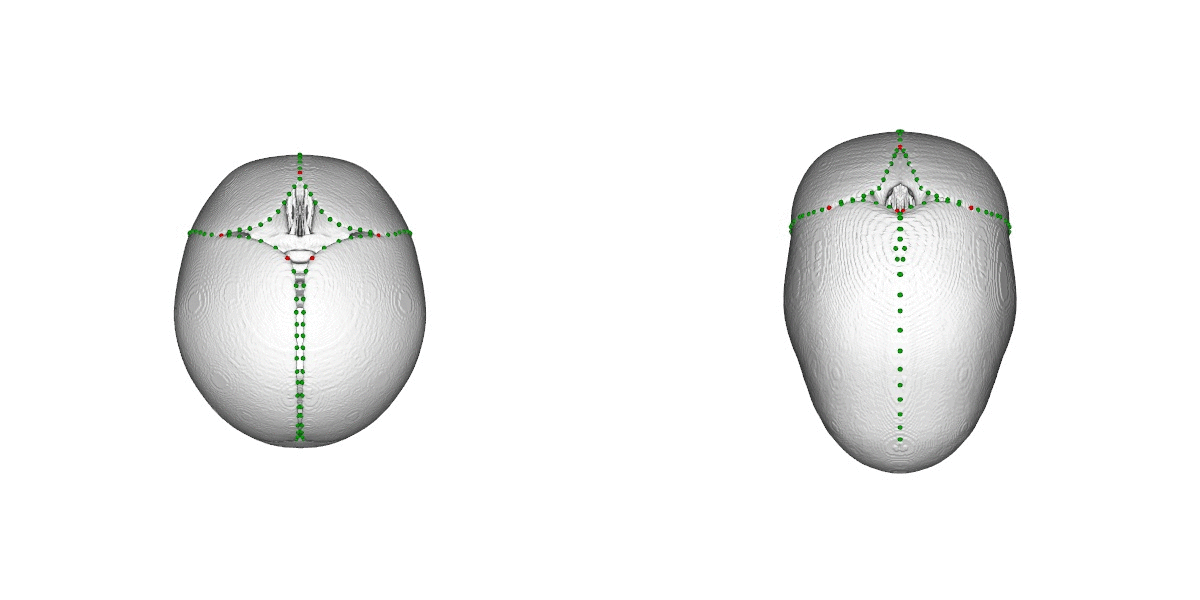

### Supplemental figure

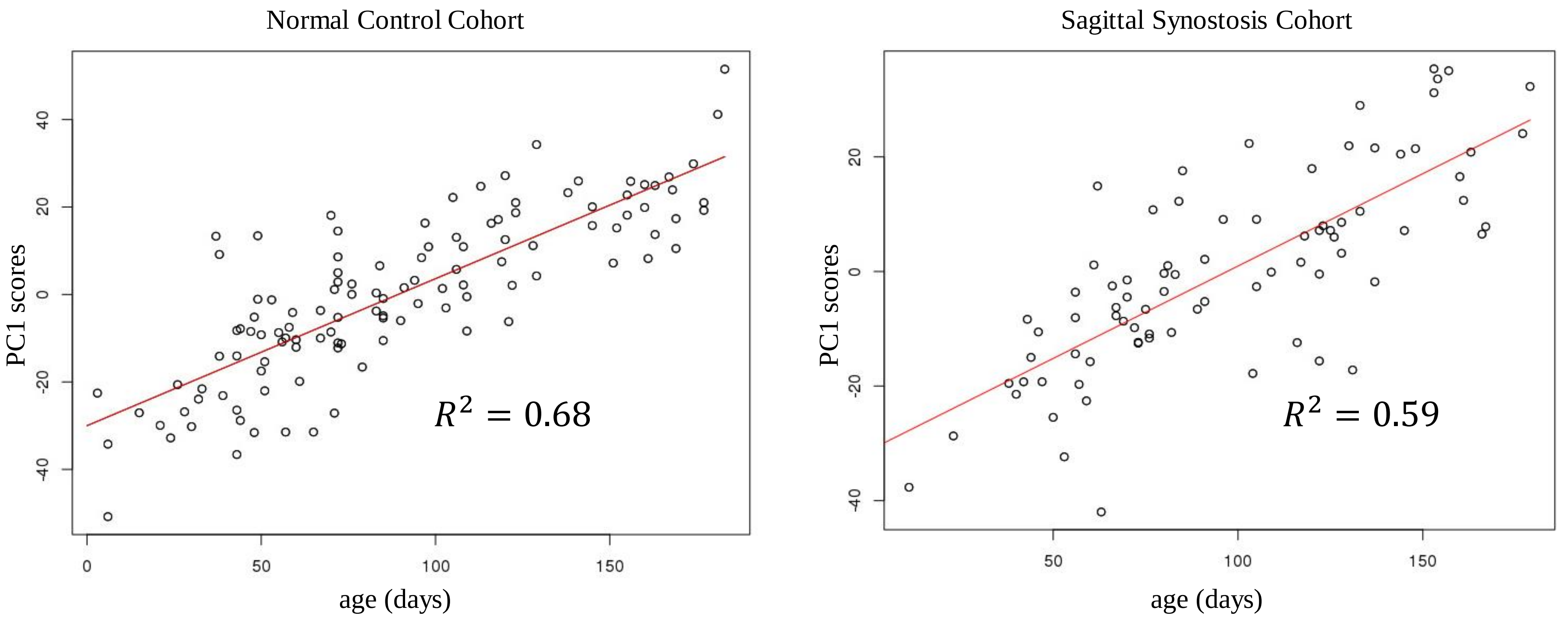

### Supplemental Figure 1

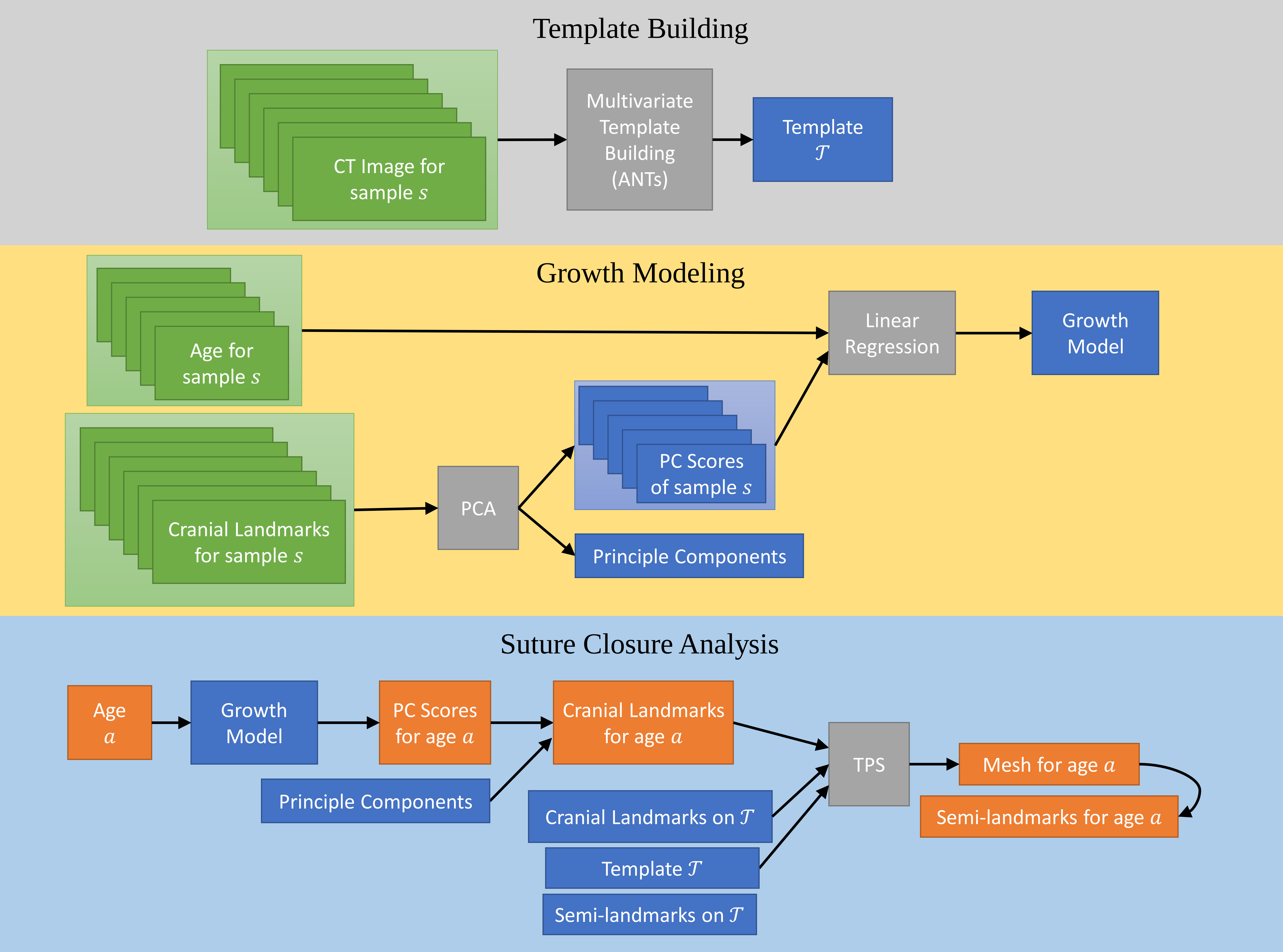
