## Appendix: Landmarks and semi-landmarks for "Cranial Bone Growth in Isolated Sagittal Craniosynostosis Compared to Normal Growth in the First Six Months of Age"

### S2 Appendix: Landmarks and Semi-landmarks

Table: Landmarks used in growth model.

| Symbol | Name | Description |
|---|---|---|
| **po** | porion | the uppermost point on the margin of the external auditory meatus |
| **zyo** | zygoorbitale | the point where the orbital rim intersects the zygomaticomaxillary suture |
| **n** | nasion | the intersection of the frontal bone and two nasal bones |
| **ns** | nasospinale | the median point of a line marking the inferior margins of the nasal apertures |
| **son** | supraorbital notch | small groove at superior and medial margin of the orbit in the frontal bone |
| **d** | dacryon | junction of the lacrimal, maxilla and frontal sutures on the medial wall of the orbit |
| **fmo** | frontomalare orbitale | the point where the frontozygomatic suture crosses the inner orbital rim |
| **fmt** | frontomalare tempotrale | the point where the frontozygomatic suture crosses the temporal line |
| **pt** | pterion | the point where the frontal, parietal, temporal, and sphenoid join together (parietal, anterior border) |
| **ast** | asterion | the point where the lambdoidal, parietomastoid and occipitomastoid sutures meet (on the occipital, anterior-most border) |
| **l** | lambda | the point of intersection of the sagittal and lambdoidal sutures in the median sagittal plane. (superior border of occipital, the mid-point closest to the suture zone) |
| **i** | inion | the highest point of the external occipital protuberance near the middle of the squamous part of occipital |
| **bp** | bregma | the point of intersection of the coronal and sagittal sutures (on parietal) |
| **bf** | bregma | the point of intersection of the coronal and sagittal sutures (on frontal) |
| **ba** | basion | the midpoint on the anterior margin of the foramen magnum |
| **o** | opisthion | the midpoint on the posterior margin of the foramen magnum |

| | | |
|---|---|---|
| **exo** | - | the medioposterior margin of the exoccipital bone |
| **bop** | - | basioccipital/basisphenoid synchondrosis posterior margin (on the anterior border of basioccipital) |
| **boa** | - | basioccipital/basisphenoid synchondrosis right anterior margin (on the posterior border of basisphenoid) |
| **se** | sella | the midline point of the dorsum of saddle-shaped depression in the body of the sphenoid bone, in the posterior view of the cranium |
| **eu** | euryon | the ectocranial point of greatest cranial breadth |
| **g** | glabella | the anterior- most midline point on the frontal bone |
| **op** | opisthocranion | the ectocranial point (at the rear of the cranium) of greatest length from glabella |
| **v** | vertex | the superior-most point of the cranium in the midsagittal contour |

The anatomical and geometric landmarks, i.e. cranial landmarks, are mostly developed for adult skull. Some landmarks cannot be identified on the young infant skulls, such as bregma (the point where metopic, sagittal and coronal sutures meet). We adapted the standard anatomical landmarks for consistency in annotation and model precision by either defining their position on specific bones (e.g. asterion is always on the occipital at the anterior-most border although, by definition, it is the point where the occipitomastoid sutures meet) or dividing them into multiple landmarks (bregma is divided into four landmarks: two on the parietal bones and two on the frontal bones with the assumption that these bones will grow to meet at bregma). Even with these attempts, the cranial landmarks do no cover the area of interest: the major sutures around frontal and parietal bones. To be able to model the growth at the sutures, we annotated them with dense semi-landmarks. Our purpose was to compare the growth in normal model with the sagittal synostosis model, hence, it was important to match the semi-landmarks approximately in both models. We divided the sutures of interest into regions based on the points of bone contact, and annotated regions with the same number of semi-landmarks in both models: region 1 (nasion to the end of metopic suture) had 12 semi-landmarks, region 2 (anterior border of fontanelle on the frontal bone) had 11 semi-landmarks, region 3 and region 4 (coronal suture) had 12 semi-landmarks each, region 5 (posterior border of fontanelle on the parietal bone) had 7 semi-landmarks, region 6 (sagittal suture) had 17 semi-landmarks and region 7 (lambdoidal suture) had 9 semi-landmarks. We placed the semi-landmarks in each region with equal spacing which meant, for example, the sagittal suture on the sagittal craniosynostosis model had semi-landmarks farther apart than those of normal model.

We also qualitatively evaluated the models by deforming the normative and sagittal craniosynostosis template meshes to landmark locations predicted at weekly intervals and creating an animation of growth. Visualizing the cranium and reviewing growth animations further showed that linear regression gives a more realistic growth model. Please see Supplementary Material S5 Figure for the growth animations.
