## Appendix: Statistical models compared for "Cranial Bone Growth in Isolated Sagittal Craniosynostosis Compared to Normal Growth in the First Six Months of Age"

### S3 Appendix: Comparison of Growth Models

We modeled growth as the landmark coordinate displacement over age in days. As regression models, we evaluated linear regression (L1), second-degree polynomial regression (L2) and locally estimated scatterplot smoothing (LOESS). As response variable (or dependent variable), we compared coordinate values after Procrustes superimposition, and as well as principle component scores from the first 20 derived from applying PCA to these Procrustes aligned coordinates. The first 20 PCs accounted for 89% and 90% of the variation in normal SCS models respectively. Our evaluations were composed of a quantitative aspect (leave-one-out-cross-validation, LOOCV) and a qualitative aspect (visual inspection of the growth on 3D meshes interpolated with thin plate spline (TPS)).

For both models, we left one sample out in the model estimation and calculated the average error as the distance between the predicted landmark coordinates and the actual landmark coordinates for this sample. By iterating over all samples, we calculated an average error for the population for each method.

Table below shows the average error for each landmark using different growth models. The differences between prediction errors are all less than 1mm and the largest prediction errors are for eurons and opisthocranion landmarks which are landmarks that defined by geometry rather than anatomy and are variable based on alignment of the skull. All the other landmarks have errors ~ < 5mm in all models.

As prediction errors were not widely different from each other, in the paper we used the simplest model: multiple-linear regression (L1) with 20 PCs which also gives us the least number of parameters.

Table: Prediction error (in mm) for landmarks using different statistical models for normal growth and sagittal craniosynostosis growth.

| | Control | | | | | | Sagittal Craniosynostosis | | | | | |
|---|---|---|---|---|---|---|---|---|---|---|---|---|
| | **LOESS** | **L1** | **L2** | **LOESS PC20** | **L1 PC20** | **L2 PC20** | **LOESS** | **L1** | **L2** | **LOESS PC20** | **L1 PC20** | **L2 PC20** |
| **euR** | 9.22 | 9.11 | 9.17 | 9.26 | 9.14 | 9.21 | 6.25 | 6.11 | 6.17 | 6.16 | 6.06 | 6.10 |
| **euL** | 8.96 | 8.94 | 8.99 | 8.97 | 8.94 | 9.00 | 5.74 | 5.70 | 5.72 | 5.74 | 5.70 | 5.72 |
| **g** | 4.42 | 4.45 | 4.48 | 4.38 | 4.41 | 4.45 | 4.70 | 4.79 | 4.81 | 4.71 | 4.80 | 4.81 |
| **v** | NA | NA | NA | NA | NA | NA | 10.92 | 10.81 | 10.94 | 10.94 | 10.83 | 10.95 |
| **op** | 8.94 | 8.86 | 8.88 | 8.97 | 8.88 | 8.89 | 7.31 | 7.40 | 7.45 | 7.28 | 7.39 | 7.43 |
| **poR** | 3.10 | 3.08 | 3.11 | 3.09 | 3.09 | 3.10 | 2.86 | 2.88 | 2.91 | 2.85 | 2.89 | 2.91 |
| **poL** | 2.89 | 2.87 | 2.90 | 2.88 | 2.88 | 2.90 | 2.48 | 2.54 | 2.56 | 2.59 | 2.65 | 2.66 |
| **zyoR** | 2.98 | 2.97 | 2.98 | 2.98 | 2.96 | 2.96 | 2.74 | 2.84 | 2.86 | 2.73 | 2.82 | 2.84 |
| **zyoL** | 2.95 | 2.95 | 2.96 | 2.95 | 2.92 | 2.93 | 2.65 | 2.73 | 2.75 | 2.66 | 2.72 | 2.74 |
| **n** | 2.62 | 2.60 | 2.61 | 2.58 | 2.60 | 2.60 | 2.25 | 2.34 | 2.35 | 2.26 | 2.35 | 2.36 |
| **ns** | 3.06 | 3.10 | 3.11 | 3.08 | 3.10 | 3.11 | 2.80 | 2.88 | 2.90 | 2.81 | 2.87 | 2.90 |
| **sonR** | 3.32 | 3.37 | 3.37 | 3.37 | 3.40 | 3.41 | 3.13 | 3.14 | 3.15 | 3.12 | 3.14 | 3.14 |
| **sonL** | 3.32 | 3.35 | 3.33 | 3.38 | 3.39 | 3.39 | 2.99 | 2.99 | 3.01 | 2.99 | 2.98 | 2.99 |
| **dR** | 2.69 | 2.64 | 2.65 | 2.62 | 2.60 | 2.61 | 2.33 | 2.34 | 2.37 | 2.25 | 2.28 | 2.30 |
| **dL** | 2.63 | 2.59 | 2.61 | 2.62 | 2.59 | 2.60 | 2.32 | 2.36 | 2.37 | 2.30 | 2.35 | 2.36 |
| **fmoR** | 2.80 | 2.82 | 2.84 | 2.79 | 2.82 | 2.83 | 2.55 | 2.62 | 2.64 | 2.50 | 2.58 | 2.59 |
| **fmoL** | 2.79 | 2.80 | 2.82 | 2.75 | 2.77 | 2.78 | 2.57 | 2.58 | 2.58 | 2.56 | 2.57 | 2.57 |
| **fmtR** | 2.81 | 2.80 | 2.81 | 2.77 | 2.79 | 2.79 | 2.71 | 2.74 | 2.73 | 2.67 | 2.70 | 2.69 |
| **fmtL** | 2.87 | 2.86 | 2.88 | 2.83 | 2.83 | 2.84 | 2.53 | 2.55 | 2.54 | 2.52 | 2.55 | 2.53 |
| **ptR** | 3.73 | 3.78 | 3.74 | 3.74 | 3.79 | 3.75 | 3.41 | 3.56 | 3.57 | 3.40 | 3.58 | 3.56 |
| **ptL** | 3.77 | 3.77 | 3.75 | 3.72 | 3.75 | 3.72 | 3.46 | 3.49 | 3.52 | 3.52 | 3.54 | 3.58 |
| **astR** | 5.13 | 5.10 | 5.11 | 5.03 | 5.01 | 5.02 | 4.71 | 4.80 | 4.78 | 4.68 | 4.77 | 4.75 |
| **astL** | 5.35 | 5.31 | 5.34 | 5.38 | 5.32 | 5.36 | 4.43 | 4.50 | 4.51 | 4.37 | 4.46 | 4.46 |
| **l** | 5.95 | 5.89 | 5.90 | 5.98 | 5.94 | 5.95 | 6.60 | 6.76 | 6.82 | 6.67 | 6.82 | 6.89 |
| **i** | 5.97 | 5.92 | 5.95 | 5.95 | 5.93 | 5.95 | NA | NA | NA | NA | NA | NA |
| **bpR** | 5.39 | 5.38 | 5.42 | 5.45 | 5.46 | 5.50 | 5.23 | 5.21 | 5.29 | 5.21 | 5.20 | 5.27 |
| **bpL** | 5.27 | 5.32 | 5.33 | 5.29 | 5.34 | 5.35 | 5.12 | 5.10 | 5.17 | 5.09 | 5.08 | 5.15 |
| **bfR** | 5.82 | 5.80 | 5.84 | 5.86 | 5.82 | 5.86 | 6.26 | 6.36 | 6.36 | 6.25 | 6.35 | 6.35 |
| **bfL** | 5.62 | 5.64 | 5.66 | 5.59 | 5.62 | 5.63 | 6.20 | 6.36 | 6.36 | 6.23 | 6.38 | 6.38 |
| **ba** | 2.45 | 2.48 | 2.47 | 2.47 | 2.48 | 2.47 | 2.26 | 2.25 | 2.23 | 2.26 | 2.27 | 2.25 |
| **o** | 3.51 | 3.49 | 3.51 | 3.57 | 3.55 | 3.59 | 3.61 | 3.57 | 3.60 | 3.66 | 3.61 | 3.63 |
| **exoR** | 3.53 | 3.56 | 3.59 | 3.53 | 3.57 | 3.60 | 3.20 | 3.21 | 3.22 | 3.21 | 3.23 | 3.24 |
| **exoL** | 3.45 | 3.50 | 3.53 | 3.48 | 3.54 | 3.56 | 3.10 | 3.15 | 3.15 | 3.11 | 3.14 | 3.14 |
| **bopR** | 2.08 | 2.12 | 2.08 | 2.10 | 2.14 | 2.10 | 2.01 | 2.01 | 1.99 | 1.97 | 1.97 | 1.94 |

| | | | | | | | | | | | | |
|---|---|---|---|---|---|---|---|---|---|---|---|---|
| **bopL** | 2.11 | 2.18 | 2.15 | 2.10 | 2.15 | 2.12 | 2.02 | 2.01 | 2.00 | 2.03 | 2.03 | 2.02 |
| **boaR** | 2.12 | 2.13 | 2.13 | 2.10 | 2.09 | 2.10 | 2.06 | 2.05 | 2.05 | 2.04 | 2.04 | 2.04 |
| **boaL** | 2.11 | 2.12 | 2.12 | 2.11 | 2.11 | 2.11 | 2.15 | 2.10 | 2.13 | 2.11 | 2.10 | 2.12 |
| **se** | 2.14 | 2.14 | 2.13 | 2.12 | 2.14 | 2.11 | 1.88 | 1.85 | 1.85 | 1.87 | 1.85 | 1.85 |
| **Avg** | **4.00** | **3.99** | **4.01** | **4.00** | **4.00** | **4.01** | **3.77** | **3.80** | **3.82** | **3.77** | **3.80** | **3.82** |
